# Interaural time difference sensitivity under binaural cochlear implant stimulation persists at high pulse rates up to 900 pps

**DOI:** 10.1101/2022.02.09.479686

**Authors:** Alexa N. Buck, Sarah Buchholz, Jan W. Schnupp, Nicole Rosskothen-Kuhl

## Abstract

**Background:** Spatial hearing remains one of the major challenges for bilateral cochlear implant (biCI) users, and early deaf patients in particular are often completely insensitive to interaural time differences (ITDs) delivered through biCIs. One popular hypothesis is that this may be due to a lack of early binaural experience. However, we have recently shown that neonatally deafened rats fitted with biCIs in adulthood quickly learn to discriminate ITDs as well as their normal hearing litter mates, and perform an order of magnitude better than human biCI users.

**Methods:** Our unique behaving biCI rat model allows us to investigate other possible limiting factors of prosthetic binaural hearing, such as the effect of stimulus pulse rate and envelope shape. Previous work has indicated that ITD sensitivity may decline substantially at the high pulse rates often used in clinical practice. We therefore measured behavioral ITD thresholds in neonatally deafened, adult implanted biCI rats to pulse trains of 50, 300, 900 and 1800 pulses per second (pps), with either rectangular or Hanning window envelopes.

**Results:** Our rats exhibited very high sensitivity to ITDs at pulse rates up to 900 pps for both envelope shapes, similar to those in common clinical use. However, ITD sensitivity declined to near zero at 1800 pps, for both Hanning and rectangular windowed pulse trains.

**Conclusions:** Current clinical cochlear implant (CI) processors are often set to pulse rates ≥900 pps, but ITD sensitivity in human CI listeners has been reported to decline sharply above ∼300 pps. Our results suggest that the relatively poor ITD sensitivity seen at >300 pps in human CI users may not reflect the hard upper limit of biCI ITD performance in the mammalian auditory pathway. Perhaps with training or better CI strategies good binaural hearing may be achievable at pulse rates high enough to allow good sampling of speech envelopes while delivering usable ITDs.

## Background

To date, cochlear implants (CIs) have been provided to over 500,000 severely hearing impaired patients across the globe (Ear Foundation, UK, 2016) and have greatly improved the quality of life of their recipients. However, substantial limitations remain in the CI hearing experience. For example, bilateral CI (biCI) users still invariably show poor performance in binaural tasks. Such tasks as sound localization and auditory scene analysis greatly benefit from the brain’s ability to process binaural spatial cues, including interaural time differences (ITDs) and interaural level differences (ILDs). While normal hearing (NH) human listeners may be able to detect ITDs as small as ∼10 µs [1], ITD sensitivity of CI patients suffering from prelingual deafness is often poor, and even rare star performers only achieve thresholds of a few hundred µs [2-9]. This poor ITD sensitivity is often hypothesized to be a result of the absence of auditory experience during an early critical period [10]. However, in [11] we used a rat model to show that, even in the absence of early auditory input, near normal ITD sensitivities can be obtained, at least for low pulse rates, when stimuli were provided with precisely synchronized CI processors. This indicates that developmental critical periods may not be the main reason for the poor ITD sensitivity seen in biCI users, and that technological shortcomings may instead be the main limiting factors.

One critical factor is likely to be the pulse rate at which CIs operate. For example, a review by [6] concludes that ITD performance of postlingually deaf biCI users tends to decline as pulse rates increase above ∼300 pulses per second (pps). Several previous studies on human listeners [12-17] as well as physiological measures of sensitivity to ITDs obtained from experimental animals [18-20] generally support this conclusion. However, accurately characterizing the dependence of ITD sensitivity on pulse rates in human listeners is marred with difficulties, and much of the literature on the ITD sensitivity of human CI listeners so far has focused only on pulse rates not exceeding 300 pps [2; 21-25]. The wide variety of patient histories and the very limited control that researchers have over variables such as the patients’ CI hardware, implantation history, stimulation parameters, experience, daily routine or availability for psychoacoustic testing introduce numerous confounds, which make it very difficult to isolate the effect of pulse rate. Indeed, previous human studies that assessed higher pulse rates reported highly variable results, but reports of any measurable ITD sensitivity at pulse rates of 300 pps or above has only been observed only very exceptionally, and only a small number of postlingually deaf CI users [12; 15; 16; 26; 27]. Thus, previous research paints a pessimistic picture of the levels of ITD sensitivity that might be achievable at pulse rates that are fast enough to allow adequate speech encoding. But patients so far have never been fitted with devices that make the delivery of abundant ITD cues a priority, nor has binaural cue sensitivity featured much in their rehabilitation. We therefore do not really know what the true potential for ITD sensitivity at high pulse rates could be under favorable conditions. To address this question, the recent development of a behavioral biCI rat model [11] is invaluable.

While concerns of species differences remain, there is more and more evidence from recent studies [28; 29] suggesting rat binaural hearing is in fact far more similar to that of humans than previously suspected. In fact, normal hearing rats exhibit behavioral high frequency (“envelope”) ITD thresholds which are very similar to those seen in humans [28], and even show very similar “precedence effects’’ as are seen in human ITD perception [29; 30]. Thus, the available psychoacoustic data show that the performance parameters of binaural hearing in rats are fundamentally similar to those of humans, and the parsimonious assumption that the functional organisation and underlying anatomy and physiology will be fundamentally similar is valid in the absence of evidence to the contrary. This makes rats a well validated model for human binaural hearing, well suited for characterizing the stimulus parameter ranges that allow high levels of ITD sensitivity with high accuracy and reproducibility.

Current CI processors generally run at fixed pulse rates between 900 and 3700 pps [31]. The rationale here is that faster temporal sampling of speech envelopes might improve speech recognition in CI users. However, [32] demonstrated that increasing pulse rates from 600 to 2400 pps resulted in little to no benefit for phoneme, word, and sentence recognition in quiet or in noise. Nevertheless, even 600 pps could be “too fast for ITD”, given the poor performance data of human CI patients at rates of 300 pps or above that we have just reviewed. Modern CI processor designers thus face conflicting demands: they must use pulse rates that are fast enough to sample the envelopes of speech and other sounds of interest with sufficient temporal resolution, yet slow enough to permit good ITD sensitivity. Judging from the current human patient literature, one may get the impression that pulse rates that allow both may not exist, in which case allowing for ITD cues will have to fall by the wayside, given that good speech encoding will clearly be the higher priority. However, neither the studies carried out so far to explore the lower limits of pulse rates required for good speech encoding, nor those exploring the upper limits of pulse rates for good ITD encoding can be considered definitive, so there is little data to guide decisions for device design or clinical practice regarding which ranges of pulse rates might be considered optimal for future biCI stimulation strategies. To start filling these important gaps in our knowledge, we conducted a series of positively reinforced two-alternative forced choice (2-AFC) ITD lateralization experiments on neonatally deafened (ND) rats which were fitted with biCIs in early adulthood, measuring their behavioral ITD thresholds with binaural pulse trains at a variety of pulse rates, and with either sharp (rectangular) or gentle (Hanning windowed) onsets and offsets. Based on previous studies in CI listeners [4; 8; 9] and electrophysiology studies in CI animals [19; 33-37] we expected to see generally better ITD sensitivities for lower pulse rates and a rapid drop off above 300 pps. In addition, we expected to observe reduced ITD lateralization performances when pulse trains were presented with gently rising slopes (Hanning windowed), rather than sharp onset rectangle windowed pulse trains, given that previous physiological and behavioral studies have consistently shown that ITD sensitivity is heavily weighted towards the onset of stimuli [29; 30]. But gently rising windows make the exact onset of the sound uncertain due to the initially very low amplitude. Our expectations regarding pulse rates and envelope shapes were only in part born out in the data, but, in striking contrast to our predictions, we found that excellent ITD sensitivity persisted at 900 pps in our early deafened biCI rats. Note that van Hoesel and colleagues [26] reported that ITD thresholds for human CI listeners were too large to measure at 800 pps. Furthermore, using Hanning as opposed to rectangular envelopes led to only modest reductions in ITD sensitivity. These results are important as they suggest that, under more optimized conditions, far better ITD sensitivity under CI stimulation might be achievable in patients than previous studies would have suggested.

## Results

Sixteen ND rats received chronic biCIs in early adulthood (postnatal weeks 10-14) and were successfully trained to localize ITD cues within 3-5 days of behavioral training (8 sessions on average with 2 sessions per day), regardless of the starting pulse rate (50, 300 or 900 pps). Figure 1 shows the experimental timeline and the number of CI animals tested per condition. For details on the behavioral training and test setup see [11] and Methods below. Representative psychometric curves for ITD lateralization with rectangular windowed pulse trains from four of these NDCI rats are shown in Figure 2. Supplementary Figure S2 shows the corresponding psychometric curves for rectangle windowed stimuli for all 16 animals tested. Uniformly good ITD sensitivity, as evidenced by steep slopes in the psychometric functions and very low error rates for absolute ITDs ≥ 80 μs, is seen at all pulse rates tested with the exception of 1800 pps. At 1800 pps the animals’ performance dropped to near-chance, with high variability in performance, as seen by the large error bars (Figs. 2, S2), even at large ITDs; whereas at lower pulse rates the same animals were performing well over 80% correct for ITDs ≥ 60 μs. In two of the CI animals shown in Figure 2, the decrease in ITD performance was accompanied by a bias for the left spout (NDCI-14 and NDCI-15).

**Figure 1:**
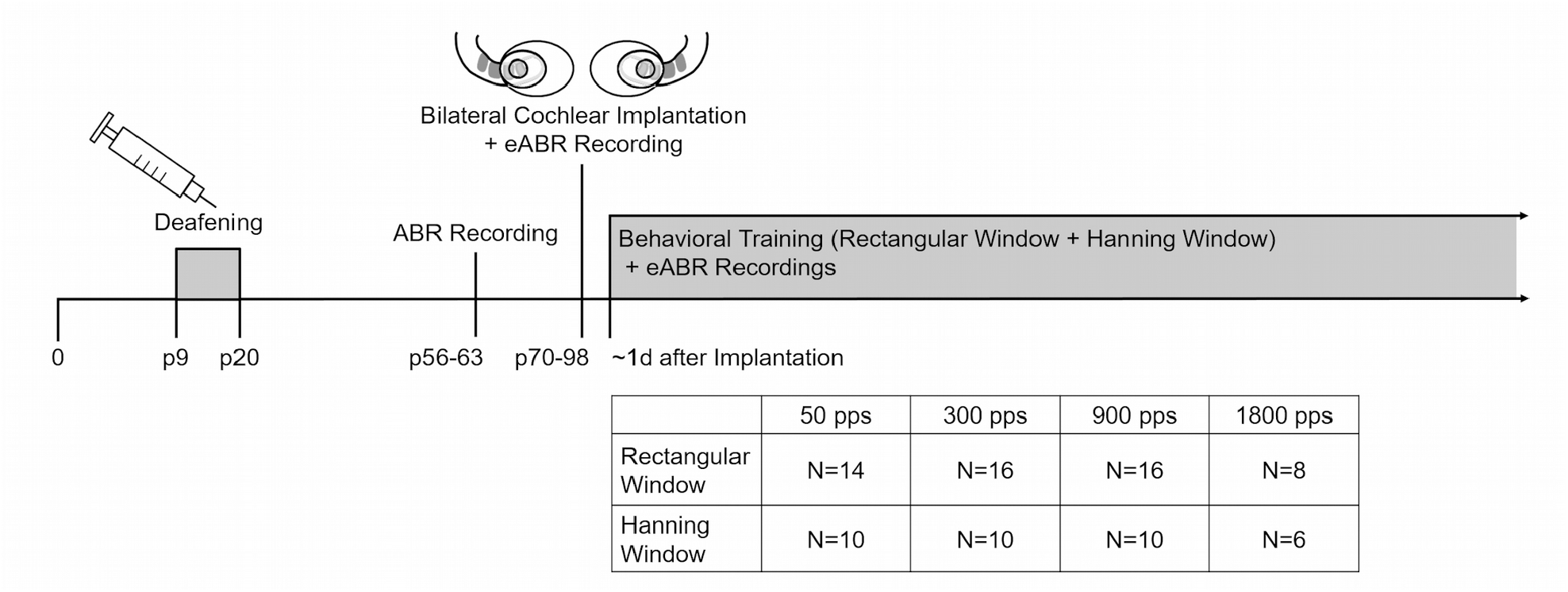
Experimental pipeline with timeline and number of animals (N) tested per condition. Abbreviations: ABR = acoustically evoked auditory brainstem response, eABR = electrically evoked auditory brainstem response, p = postnatal day, pps = pulses per second, d= day.

**Figure 2:**
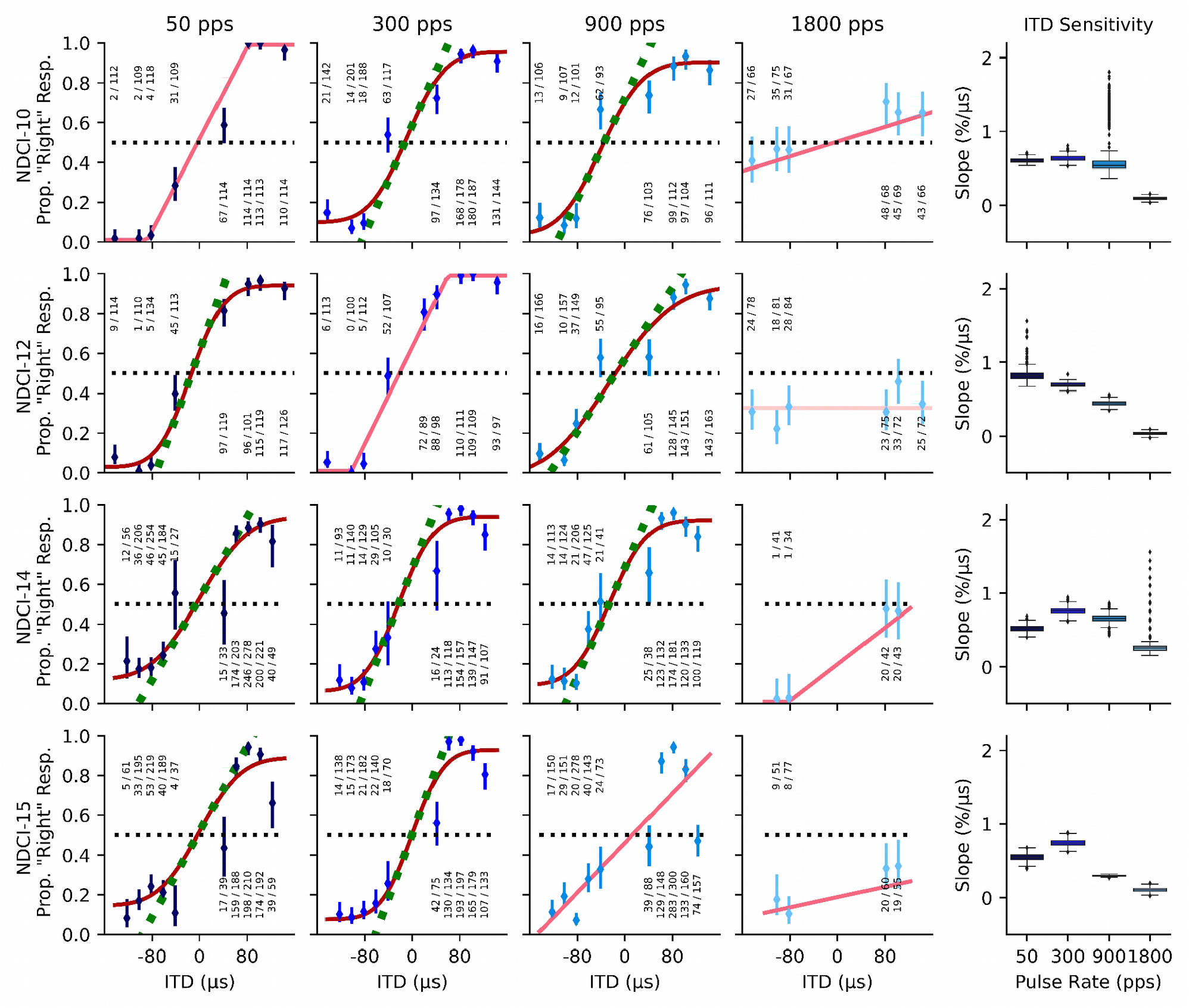
Example psychometric curves for four neonatally deafened, cochlear implanted (NDCI) rats localizing rectangular windowed pulse trains by interaural time difference (ITD). The first four columns represent a different pulse rate from left to right 50, 300, 900, and 1800 pulses per second (pps), each indicated with a different shade of blue. The y-coordinates reflect the proportion of trials during which the animal responded on the “right” hand spout (Prop. “Right” Resp.). The x-axis shows the tested ITD values from – 150 to + 150 µs. Negative ITD values indicate left ear leading ITDs. Annotations above or below each marker indicate the number of trials the animal chose the right hand side spout over the total number of trials for the given ITD value on the x-axis. Dark red curves show sigmoidal fits to the responses, bright red lines are linear fits with bounds, light pink are null model psychometric curve fits (see “Data analysis” above for details on the model fits). Green dashed lines show the slopes of psychometric curves at ITD=0 µs. The slopes serve to quantify the behavioral sensitivity of the animal to ITDs. The fifth column summarizes the ITD sensitivity across the different pps per animal as a function of slope. Each row shows the responses for a given animal. Psychometrics of all CI animals are shown in Figure S2.

In addition, 10 out of 16 animals were tested for ITD sensitivity to 50, 300 and 900 pps (N=10), as well as at 1800 pps (N=6), pulse trains which were amplitude modulated with a slow rising and falling Hanning window (see Figs. 1 and S3 for details). Again, all animals showed good to excellent ITD sensitivity with steep psychometric slopes at all pulse rates except 1800 pps (Fig. S3). At this pulse rate, the psychometric functions were completely flat (slopes not statistically different from zero) for 5 out of 6 animals tested, as shown in the fourth column of Figures 3 and S3. The slopes of the psychometric functions for Hanning windowed pulse trains were generally found to be a little shallower than those from the sharp onset rectangular windowed stimuli shown in Figures 2 and S2.

**Figure 3:**
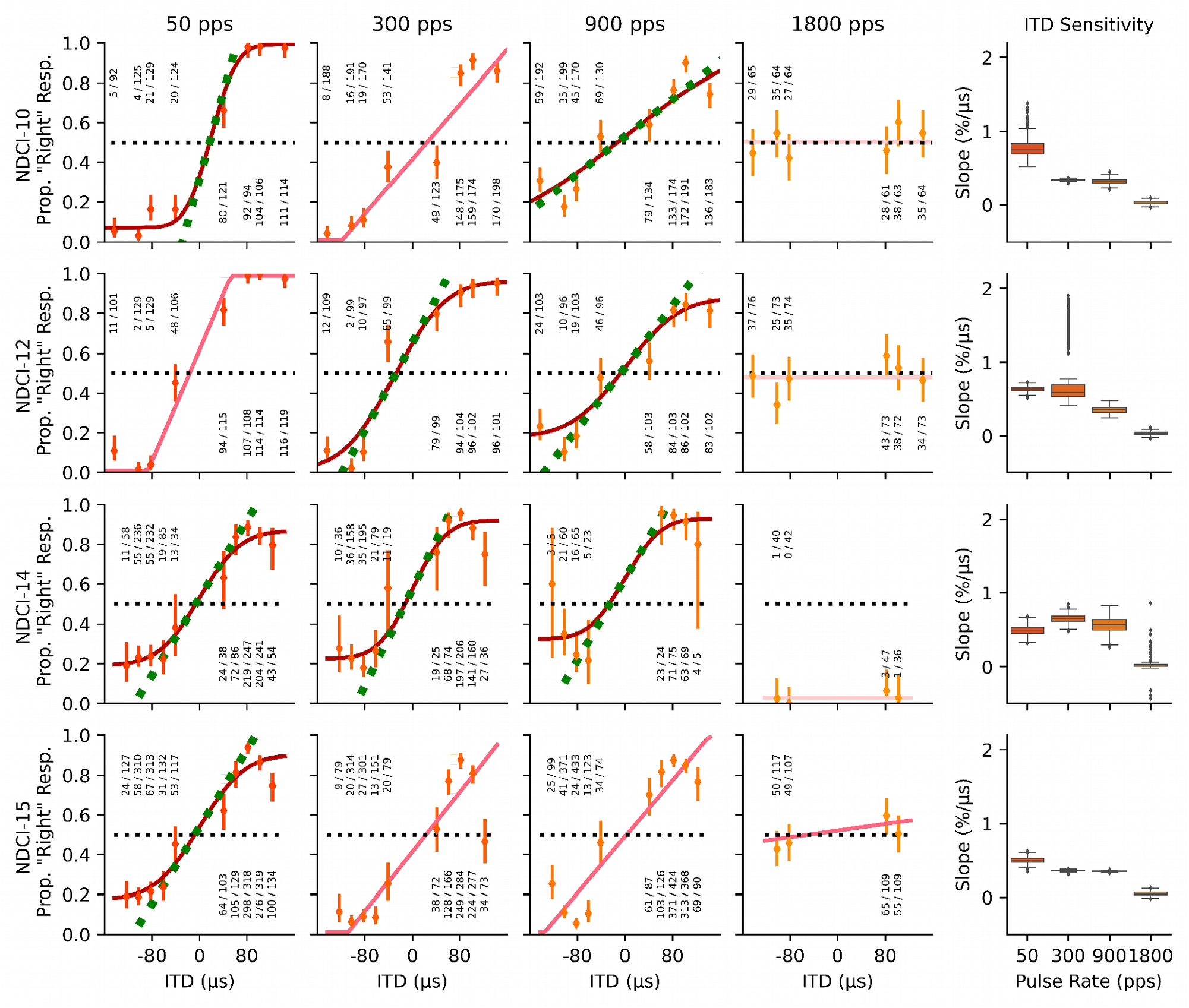
Psychometric curves for the same four example neonatally deafened, cochlear implanted (NDCI) rats as shown in Figure 2 localizing Hanning windowed pulse trains with varying interaural time differences (ITDs). Details are as for Figure 2. Psychometrics of all CI animals tested on Hanning windowed stimuli are shown in Figure S3.

To quantify the effect of pulse rate and pulse train envelope on ITD discrimination performance in biCI animals, we quantified behavioral sensitivity by the slope of the psychometric curve around the ITD=0 µs midline. We then used bootstrap tests (see Methods for details) to estimate the distributions and confidence limits of these sensitivity values for all animals at each pulse rate and envelope type, and to estimate the probability that the observed differences in mean ITD sensitivity across animals as a function of either pulse rate or envelope type were larger than expected by chance. Figure 4 shows the bootstrapped mean distributions of ITD sensitivities as a function of stimulus window type and pulse rate, averaged over all animals for which behavioral data at the relevant combination of stimulus parameters was available. The p-values for the null hypothesis that ITD sensitivity did not differ for a given pair of stimulus parameter combinations were Bonferroni corrected to account for multiple comparisons and are listed in Tables 1, 2, and 3. ITD sensitivities were usually higher with rectangular than Hanning windowed envelopes, and that effect was statistically significant at 300 and 900 pps. ITD sensitivity also tended to decline, often significantly, with increasing pulse rate, but good ITD sensitivity was nevertheless common at pulse rates as high as 900 pps, even with Hanning windowed stimuli. In contrast, at 1800 pps, ITD sensitivity was consistently very poor, irrespective of whether rectangular or Hanning pulse train envelopes were used. Psychometric slopes are not very commonly used as metrics for sensitivity. To help readers put the observed slope values into perspective and compare them against ITD discrimination thresholds reported in other previous studies, we also plot 75% correct discrimination thresholds estimated from the psychometric, in µs, on the right y-axis of Figure 4. Some authors, including [38] and [39], refer to 75% discrimination thresholds as the “just-noticable difference” (JND). Note, however, that our animals were only tested on a lateralization task, not a discrimination task. Task details can influence performance, and different methods used for estimating thresholds from data can also lead to somewhat different values, so readers may wish to use some discretion when comparing our JND scores against those reported elsewhere. Nevertheless, these values should allow at least a rough comparison. Table 4 gives the mean of the 75% discrimination thresholds, or JNDs, as estimated from the mean slopes of the psychometric curves (see Methods), for each of the eight different stimulation conditions. Overall, the smallest thresholds were obtained for pulse rates of 50 and 300 pps, and rectangular window stimuli yielded lower thresholds. However, even at a clinically relevant pulse rate of 900 pps, estimated JNDs were in the order of 50 - 60 µs, which is very much lower than the ∼200 to well above 1000 ITD µs discrimination thresholds reported in the literature for early deafened biCI patients [4; 6; 8; 9; 21; 26; 40; 41]. Note that the values shown in Table 4 for 1800 pps represent gross, linear extrapolations of very small psychometric slopes which cannot be estimated with great precision and which lie well beyond the range of ITDs tested. Thus, the mean threshold values shown for 50, 300, and 900 pps can be considered fairly reliable and accurate (a simple visual comparison against the psychometric functions shown in Figs. 2, 3, S2, and S3 confirms that they must be “in the right ballpark”), but those at 1800 pps could be off by several 100%, given that most animals never approached 75% correct for any of the 1800 pps stimuli tested, and the data therefore very poorly constrain these estimates.

**Table 1:** Bonferroni corrected p-values for the null hypothesis that mean ITD sensitivities do not vary as a function of pulse rate for rectangular windowed stimuli with pps values shown in the row and column headings. Significant p-values (<0.05) are shown in bold.

| Rectangle | 50 pps | 300 pps | 900 pps |
| --- | --- | --- | --- |
| 300 pps | 0.546 |  |  |
| 900 pps | 0.071 | **0.001** |  |
| 1800 pps | **<0.001** | **<0.001** | **<0.001** |

**Table 2:** p-values as in Table 1, but for Hanning windowed stimuli.

| Hanning | 50 pps | 300 pps | 900 pps |
| --- | --- | --- | --- |
| 300 pps | **0.01** |  |  |
| 900 pps | **<0.001** | 1 |  |
| 1800 pps | **<0.001** | **<0.001** | **<0.001** |

**Table 3:** Bonferroni corrected p-values for the null hypothesis that mean ITD sensitivities do not vary as a function of envelope shape, rectangular or Hanning windowed stimuli, when tested at the pps values shown in the column headings. Significant p-values (<0.05) are shown in bold.

|  | at 50 pps | at 300 pps | at 900 pps | at 1800 pps |
| --- | --- | --- | --- | --- |
| Hanning vs. Rectangle | >0.999 | **<0.001** | **0.028** | 0.062 |

**Table 4:** Mean 75% ITD discrimination thresholds in µs, estimated from the slopes of the psychometric curves (see Methods) and averaged over all animals (N) tested at the corresponding stimulus condition. Note that the values shown for 1800 pps (marked †) are estimated by extrapolation well beyond the tested range. While we can be confident that thresholds at 1800 pps will be much larger than thresholds for ≤ 900 pps, our data does not permit an accurate estimation of 75% thresholds for 1800 pps, so the values shown here should not be taken at face value.

| 75% threshold | at 50 pps | at 300 pps | at 900 pps | at 1800 pps |
| --- | --- | --- | --- | --- |
| Rectangle | 43.2 (N=14) | 34.6 (N=16) | 50.5 (N=16) | 401.5<sup>†</sup> (N=8) |
| Hanning | 40.5 (N=10) | 61.8 (N=10) | 62.0 (N=10) | 3575.8<sup>†</sup> (N=6) |

**Figure 4:**
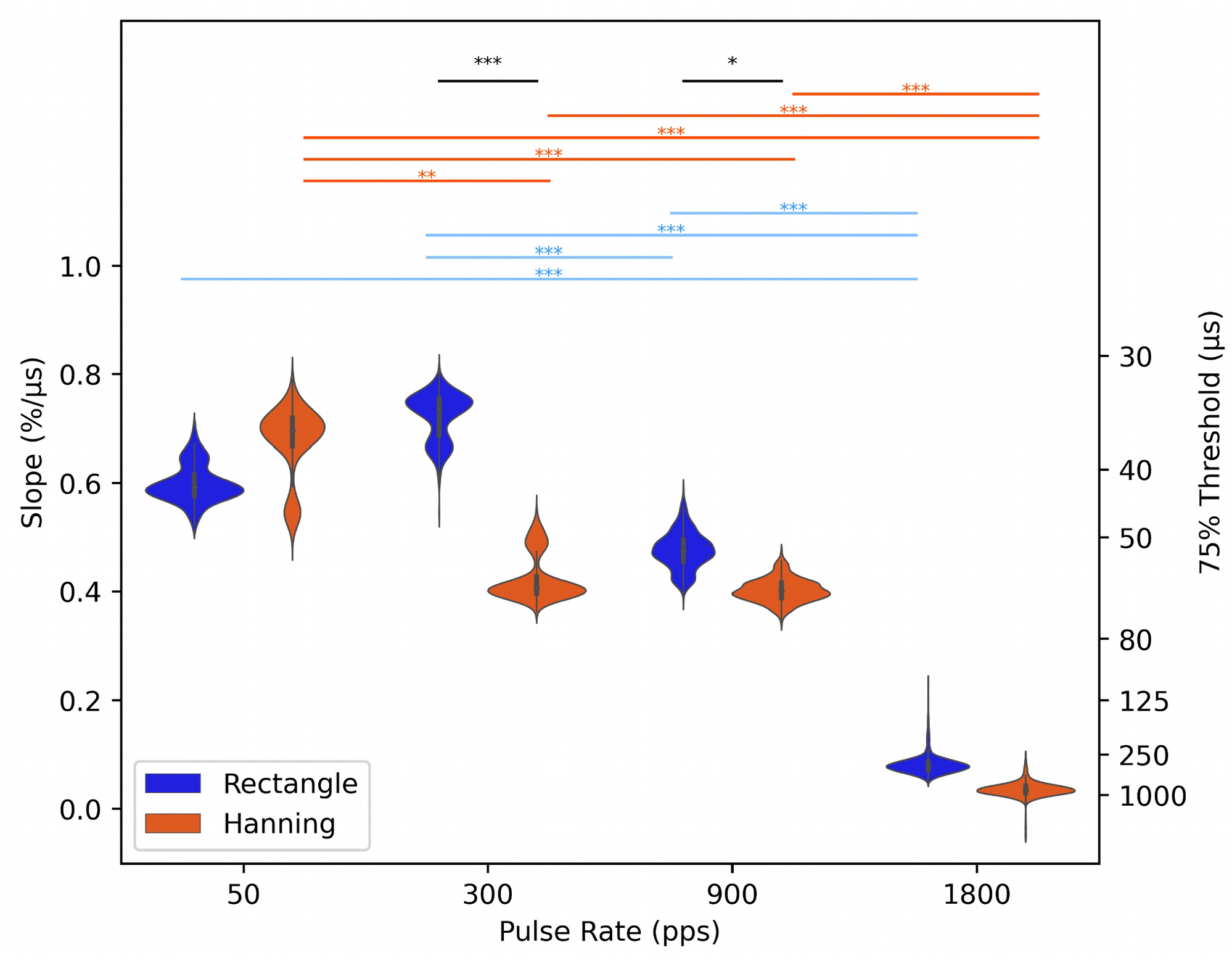
Violin plots showing the distribution of the mean group effects of ITD sensitivity for each pulse rate following permutation for rectangular (blue) and Hanning (orange) windowed stimuli. The left axis shows the ITD sensitivity as a function of the psychometric slopes, and the right y-axis the corresponding 75% ITD discrimination threshold estimate. Significant group statistics for the slopes comparing pulse rates are shown above violin plots for rectangular (blue) and Hanning (orange) windowed data. Significant differences between the two envelopes are shown in black. *** = (p<0.001), **= (0.001≤p<0.01), * = (0.01≤p<0.05).

## Discussion

### ITD lateralization in neonatally deafened CI rats is much better than expected at pulse rates up to 900 pps

In this study, we have demonstrated remarkably good ITD sensitivity despite early onset deafness under bilateral electric stimulation, including at electrical pulse rates in the range of those commonly used in current clinical practice for the purpose of accurately encoding speech envelope cues. This is directly apparent from the data even without statistical analysis. The psychometric curves shown in Figures 2, 3, S2, and S3 show that, even with Hanning windowed stimuli, all of our biCI rats routinely performed above 80% correct when presented with ITDs of ∼80 µs at pulse rates of 900 pps. Their performance is thus very substantially better than that seen in human CI listeners, who frequently show no measurable ITD sensitivity at 300 pps or above, and who, even at lower pps, fail to reach thresholds below several hundred µs [2; 16; 26]. To overcome the well documented limitations in ITD sensitivity in biCI listeners, new strategies have been tried in studies on humans [42-44] and animals [45]. For example, configurations with mixed stimulation rates, in which one low rate electrode was included among an array of higher rate electrodes, inducing ITD sensitivity similar to that seen with configurations comprising only low rates [44]. In addition, stimulation strategies have been developed that allow the encoding of the fine structure of sound. In [42], 50% of the listeners using such a fine structure strategy showed an ITD sensitivity with a mean threshold at 330 µs, while listeners without fine structure strategy failed to detect ITDs > 1000 µs. Another proposed strategy involved combining both high and low pulse rates together. Srinivasan and colleagues [43] reported that the introduction of an additional slower pulse train overlying a faster pulse train, but with a small offset, improves ITD sensitivity beyond what might be expected from a simple, equivalent increase in stimulus amplitude. In biCI rabbits, Buechel and colleagues [45] have shown that adding extra pulses to periodic pulse trains of high pulse rates introduces short inter-pulse intervals resulting in better ITD coding of auditory midbrain neurons. These new stimulation strategies are ingenious, but even with the improvements that they may have induced, the ITD discrimination of the human CI listeners tested still falls short of the performance that our early deafened rats were able to achieve at high pulse rates of up to 900 pps and without special strategies. This suggests that the factors limiting ITD perception in human biCI users may originate from factors other than carrier rates or missing hearing experience during early development.

Interestingly, the CI rats in this study even outperformed their normally hearing, acoustically stimulated peers, whose ITD sensitivity, as reported in [28], drops off dramatically as acoustic pulse rates increase from 300 to 900 pps. This is an intriguing observation. A priori, one might expect that electrical stimulation to lead to very precisely stimulus locked and highly synchronized temporal firing in the auditory nerve fiber array as it bypasses sources of noise and temporal jitter in the auditory periphery, cochlear hair cells, or the hair cell synapses. This, in turn, ought to allow for better, rather than worse, temporal encoding and processing than can be achieved with acoustic stimuli. It is therefore disappointing that temporal processing in CI patients has so far been found to be generally poorer than that reported for normally hearing individuals, but the data presented here indicate that this need not necessarily always be the case.

Human CI users who do exhibit at least some ITD sensitivity, mostly postlingually deaf, tend to show a rapid drop-off in ITD lateralization performance with pulse rates at or above 300-800 pps [6], and their ability to use ITD cues is also lower in stimuli with speech waveform envelopes [46; 47] when compared to pulse trains with sudden, rectangular onset. Our data show similar trends, but they do strongly suggest that the cut-offs observed in human studies so far likely significantly underestimate the performance that the mammalian auditory pathway is capable of if binaural stimulation conditions are optimized. While there is evidence that CI users may be able to use ITDs carried on slow rising envelopes, human studies demonstrated that this is only possible when the shape of the envelope is peaked [26; 46; 48; 49]. This strongly suggests that a speech envelope, or any slow rising envelope for that matter (such as a Hanning windowed pulse train), does not contain the temporal features needed for effective encoding of ITD information when delivered by current clinical speech processors.

The fact that ITD discrimination is generally better for stimuli with rapid onsets rather than slowly rising envelopes is perhaps unsurprising given that, for normal hearing subjects, onset ITDs are normally the most salient [29; 30]. Electrophysiological studies of ITD sensitivity have similarly demonstrated the importance of onset cues in ITD sensitivity [29; 50; 51]. For example, in the inferior colliculus, ITD sensitivity may drop by 24-48% when onset responses are excluded from the analysis [50]. In addition, physiological ITD sensitivity under biCI stimulation has been shown to decrease with increasing pulse rates in studies when the onset cues were both included [19] or excluded [18; 20; 34; 35]. For example, [19] identified the best ITD sensitivity in inferior colliculus for pulse rates around 80-160 pps followed by a degraded sensitivity for higher pulse rates. However, in contrast to lower pulse rates, which are expected to improve ITD sensitivity [15; 16; 19; 52], there is evidence that temporal encoding is only improved at higher electrical pulse rates [53; 54] which again is interesting based on the evidence that ITD processing and temporal acuity may have similar mechanics [53; 55].

Our behavioural results in biCI animals presented here thus show similar trends to those seen in previous electrophysiological work, namely that ITD sensitivity may decline if stimulus envelopes are lacking pronounced onsets or if pulse rates are high [19; 34; 35; 50], but to the best of our knowledge we are the first to demonstrate this behaviourally.

### Does the place of stimulation matter?

One concern that may slow the development of new CI coding schemes that would be more appropriate for the delivery of ITDs is the question how important it is to target specifically low frequency parts of the auditory nerve. Classic views of binaural processing, often referred to as the “duplex theory”, posit that ITDs are the predominant binaural cue for low frequency sounds [56]. Given that human intracochlear electrodes implanted with current tools and surgical techniques usually fail to reach the apical, most low-frequency end of the cochlea [57], one might expect that this could reduce the potential for ITD sensitivity in biCI patients overall. Furthermore, it might be expected that the best ITD sensitivity could be observed when the apical-most electrodes are stimulated. Such considerations have led some researchers to use modiolar, auditory nerve penetrating electrodes in an animal model which easily reach the most apical, as well as more basal fibers. At least for monaural temporal pattern stimuli, low frequency pathways indeed seem to provide somewhat better temporal acuity than higher frequency fibres are stimulated with the same types of electrodes [58; 59]. Nevertheless, difficulties in stimulating specifically low frequency fibers do not seem to be a significant limiting factor for the delivery of ITDs over biCIs. It has been shown a long time ago that normal hearing listeners can exhibit thresholds as low as a few tens of μs for so-called “envelope ITDs” of high frequency carriers [60], and there is a growing awareness among researchers that high-frequency-biased brainstem nuclei, like the lateral superior olive, can exhibit pronounced ITD sensitivity when presented with brief transient of amplitude modulated stimuli [61]. Thus, classic “duplex theory” notions that ITD processing is relevant only for low frequency channels, or that, by implication, animals with poor low frequency hearing are fundamentally unsuitable as animal models for binaural hearing, are clearly a gross oversimplification. High frequency parts of the auditory pathway are clearly capable of processing ITDs of pulsatile, transient or amplitude modulated stimuli with high acuity, and that seems to be true in all mammals tested so far. Similarly, studies conducted so far on human biCI patients suggest that delivering ITDs over a wider part of the cochlea may be a more effective strategy than targeting the low frequency part of the cochlea specifically [32; 44]. In fact, studies comparing ITD sensitivity over the range of mid to high frequency parts of the cochlea, a range normally accessible with current clinical devices, have not observed a clear advantage of the lower frequency channels [4; 62]. The experiment reported here was not designed to examine possible place-of-stimulation effects, but it is noteworthy that here, like in our previous studies [11; 50], we found very high sensitivity to ITDs in the rat, even though we stimulated mid frequency parts of the cochlea. These regions are not part of what traditionally would be considered the low frequency ITD pathway, but which is equivalent to parts of the human cochlea that are routinely covered by conventional CI electrode arrays. Our results thus add to the growing evidence that delivering ITDs at any part of the cochlea can be beneficial, and that future binaural strategies should prioritize delivering ITDs over wide ranges of the cochlea, rather than attempting to target specifically the low frequency, apical regions.

### Why is ITD perception in ND biCI rats better than in human biCI users?

Why exactly our CI animals perform so much better than current human CI users we do not yet know, but we believe that the most likely cause for the difference in performance is that our animals received informative ITDs with microsecond precise relative timing of stimulus pulses delivered to each ear from the very onset of their CI stimulation. In contrast, current clinical processors do not normally deliver informative ITDs in the timing of the stimulus pulses, so most of the time biCI patients will receive ITD cues in the envelopes of the CI pulse trains only, and the extent to which the CI stimulated auditory pathway is capable of making good use of envelope-only ITDs is highly uncertain. Given that the electrical stimulation provided with contemporary CI processors provides such poor temporal fine structure information, it should not come as a surprise that prelingually deaf CI users often fail to develop any usable sensitivity to ITDs [8; 22].

Why our rats exhibit much better ITD performance than that previously reported in biCI users is not yet clear, but possible explanations, some of which we have touched on above, include species differences, differences in the nature of the stimulation and/or training effects. It is difficult to completely rule out species differences, but this explanation nevertheless seems unlikely given the numerous studies that indicate that rats are a good animal model for human cochlear implantation [11; 63-69] and binaural hearing [11; 28; 29]. We were able to demonstrate that rats have excellent ITD sensitivity with thresholds very similar to those seen in other mammals [28], and exhibit a similar precedence effect, with strikingly similar temporal weighting functions to those seen in humans [29]. Although there is evidence for anatomical differences of auditory nuclei between rats and mammals with well-developed low frequency hearing, including humans, for example for the medial superior olive [70], the best psychoacoustic data available indicate that the psychoacoustic performance parameters of binaural hearing in rats are fundamentally similar to those of humans. Consequently the assumption must be that the functional organization of the underlying auditory networks will therefore also be fundamentally similar. For example, despite differences in size of MSO between rodents with limited low frequency (e.g mice or rats) versus good low frequency hearing comparable to humans (e.g gerbils), shared anatomical, morphological and physiological properties have been demonstrated across these species [71]. These common properties include spiking pattern, bilateral inhibitory and excitatory tuning as well as improved responses with bilateral auditory stimulation, all of which are believed to play a role in ITD encoding [71-73]. Furthermore, comparable fine structure ITD sensitivity is found in the lower frequency region of the LSO and the MSO [74]. We believe that the excellent behavioral sensitivity to the ITD of pulsatile stimuli we observe here is likely mostly due to ITD processing in the LSO, and this seems to be a widespread phenomenon that is, as far as we know, implemented in similar ways across much of the mammalian kingdom [61].

Species differences are thus an unlikely explanation for the surprisingly good ITD discrimination performance of our rats, and procedural differences seem much more likely to be responsible. As already noted, our setup always provides informative ITDs with microsecond precision from the first stimulation, but processors in current clinical use do not. Standard clinical devices are typically based on two separate monaural processors with asynchronous clock times, and the timing of pulses is for the most part unrelated to the temporal fine structure of the input at each ear. Many human CI psychoacoustic studies are performed with research interfaces capable of more precise binaural synchronization, but the vast majority of the auditory experience of the participants nevertheless comes from asynchronous clinical CI speech processors, which deliver interaural pulse timing patterns that are uninformative and potentially misleading. The differences in ITD sensitivity could simply be driven by the fact that ITDs are always useful and reliably delivered to our biCI rats, but are mostly useless and misleading for the typical human biCI users, so that their auditory pathways may be incentivized to become insensitive to these cues. This hypothesis is supported by a recent electrophysiological study by [75], which has demonstrated that animals that experienced reliable ITD information through “ITD-aware sound processors’’ have a higher neuronal ITD sensitivity than biCI animals supplied with standard clinical processors.

It has long been thought that the absence of binaural experience during early life may preclude the development of normal ITD sensitivity in biCI listeners [22; 76]. However, in [11] we demonstrate that a long period of hearing deprivation from ∼p15 until early adulthood in our rats (∼p70-98) does not impair the ability to use ITDs. In fact, the ITD sensitivity of neonatally deafened, adult implanted rats was found to be comparable to that of normally hearing, acoustically stimulated rats [28]. ITD sensitivity thus does not appear to have a critical period. Providing auditory input with consistently informative ITDs from the onset of stimulation may play a much more important role in the development of ITD sensitivity than ensuring that this initial stimulation happens very early. Unlike typical early deaf human biCI users, our NDCI animals were able to lateralize ITDs with thresholds around 50 µs that appear no worse than those seen in their normally hearing siblings [28]. The rats in these studies spent their entire lives before puberty without any auditory input, but, unlike human CI users, once they were implanted they were also never exposed to prolonged periods of stimulation with the uninformative and potentially misleading pulse timing ITDs that wearers of bilateral clinical speech processors will experience constantly. Note that recent studies have demonstrated that lateralization training can lead to a reweighting of binaural cues in normal hearing listeners [77; 78] and biCI users [79], such that the listeners rely more or less heavily on ILDs or ITDs, respectively, depending on which cue is more reliable or informative. The auditory pathway thus seems clearly capable of desensitizing to ITD cues if provided with prolonged stimulation that is bereft of informative ITDs.

Finally, one needs to consider the possibility that training effects could play a role. For our biCI rats, lateralizing binaural pulse trains with precise ITDs but no other noteworthy features day after day constituted the entirety of their auditory experience, and they would spend approximately 10 hours a week, for several months, practicing the lateralization of these stimuli, with instant positive reinforcement for correct performance. In contrast, human biCI users spend most of their auditory experience trying to make sense of the cacophony of everyday life, and little, if any, of their time and attention is dedicated to honing their ITD discrimination skills. In this context it is of interest to note [80] report that just 32 hours of periodicity pitch training could lead to very substantial improvements in CI pulse rate discrimination. If monaural temporal processing in human CI users can improve with training, then the same may also be true for binaural temporal processing, as rate discrimination and ITD lateralization may share some common physiological mechanisms [81]. Generally, psychoacoustic limits observed in largely untrained CI listeners may substantially underestimate the physiological limits that the CI stimulated auditory pathway is in principle capable of. In the light of our findings here we therefore consider it highly likely that dramatic improvements in binaural hearing of human CI users ought to be achievable.

## Conclusion

In an animal model of early onset deafness and biCI stimulation, we have demonstrated that excellent behavioral ITD sensitivity can be observed even at pulse rates as high as 900 pps. ITD thresholds well below 100 μs were consistently observed for 900 pps stimuli with and without slowly rising envelopes, but sharp onset stimulus envelopes and lower pulse rates can further improve ITD sensitivity. Thus, under appropriate stimulation conditions, neither high carrier rates nor slow rising envelopes preclude the ability of biCI users to make use of ITD cues. This gives cause for optimism that it may well be possible to develop binaural CI processing strategies capable of delivering highly informative ITD cues without having to compromise on speech envelope sampling rates.

## Methods

All procedures involving experimental animals reported here were approved by the Department of Health of Hong Kong (#16-52 to 16-55; 18-81 to 18-83; 20-143 to 20-144 DH/HA&P/8/2/5) and/or the Regierungspräsidium Freiburg (#35-9185.81/G-17/124), as well as by the appropriate local ethical review committee. Sixteen female Wistar rats were used in the work presented here. All rats underwent neonatal deafening, acoustic and electric evoked auditory brainstem recordings, bilateral cochlear implantation, and behavioral training as described in [11] and briefly below.

### Neonatal Deafening

All animals were neonatally deafened as described in [64]. Briefly animals received daily kanamycin injections from postnatal day 9 to 20 inclusively. This is known to cause widespread death of inner and outer hair cells while keeping the number of spiral ganglion cells comparable to that in untreated control rats [82-84]. We subsequently verified that this had resulted in profound hearing loss (>90 dB) by the loss of Preyer’s reflex [85], the absence of hair cells in the cochlea, the absence of auditory brainstem responses (ABRs) to broadband click stimuli as well as pure tones (at 500, 1000, 2000, and 8000 Hz).

### Cochlear implantation

The animals were raised to young adulthood (postnatal weeks 10-14) in normal housing environment after which the rats were implanted simultaneously with bilateral CIs either from PEIRA (animal arrays ST08.45, ST05.45 or ST03.45, Cochlear Ltd, Peira, Beerse, Belgium) or from MED-EL (3-patch animal arrays, Medical Electronics, Innsbruck, Austria). Electrodes were inserted through a cochleostomy window over the middle turn of the cochlea corresponding to the 8-16 kHz region.

### Electric Stimulation

The electrical stimuli used to examine the animals’ electrically evoked auditory brainstem responses (eABRs), and the behavioral ITD sensitivity were generated using a Tucker-Davis Technology (TDT, Alachua, FL) IZ2MH programmable constant current stimulator at a sample rate of 48,828.125 Hz. The most apical ring of the CI electrode served as stimulating electrode, the next ring as ground electrode. All electrical intracochlear stimulation used biphasic current pulses similar to those used in clinical devices (duty cycle: 40.96 µs positive, 40.96 µs at zero, 40.96 µs negative), with peak amplitudes of up to 300 μA, depending on eABR thresholds and informally assessed behavioral comfort levels (rats will scratch their ears frequently, startle or show other signs of discomfort if stimuli are too intense). For behavioral training, we stimulated all neonatally deafened, cochlear implanted (NDCI) rats 2-6 dB above these thresholds depending on the pulse rate. Behavioral stimuli from the TDT IZ2MH were delivered directly to the animal through a custom built head connector that was connected and disconnected before and after each training session. As before, animals received binaurally synchronized input from the first stimulation. For full details on the electric stimuli and stimulation setup see [11].

### Psychoacoustic Training and Testing

The ability to discriminate ITDs of CI stimuli was tested in a 2-AFC ITD lateralization task in our custom made behavior setup as described in [11]. Briefly, this setup consists of three spouts from which the animal could receive a water reward. A light would indicate the start of the trial at which point the animal was trained to lick the centre spout which would trigger bilateral electric stimulation. The animal would then need to make a behavioral choice to lick the left or right spout and would be rewarded with water or punished with a time out and flashing light depending on whether the response corresponded to the ITD presented or not. Training and testing for the 16 animals was done in a pseudo random order following implantation (Fig. 1). Animals were tested on four pulse rates, 50, 300, 900, and 1800 pps, with rectangle (N=16) and Hanning (N=10) windowed pulse trains. For this experiment, both individual pulses and envelopes carried the same ITD information. As in [11], the ITDs were chosen from the range -150 to +150 μs, where negative represents a left leading and positive a right leading ITD. This range covers 125% of the animal’s physiological range, which is between -120 and +120 μs [86]. Each pulse train had a duration of 200 ms, and animals were stimulated at a maximum of 6 dB above their eABR thresholds (see [11] for details). Well differentiated eABRs of an example rat are shown in Figure S1 directly after implantation, 6 weeks, and 6 months post implantation. The starting frequency of the first training was assigned pseudo randomly to rule out any effects from the order in which they were trained and tested. Hanning windowed pulse trains consisted of a raised cosine waveform with a 100 ms rising and falling phase, respectively (Fig. 5).

**Figure 5:**
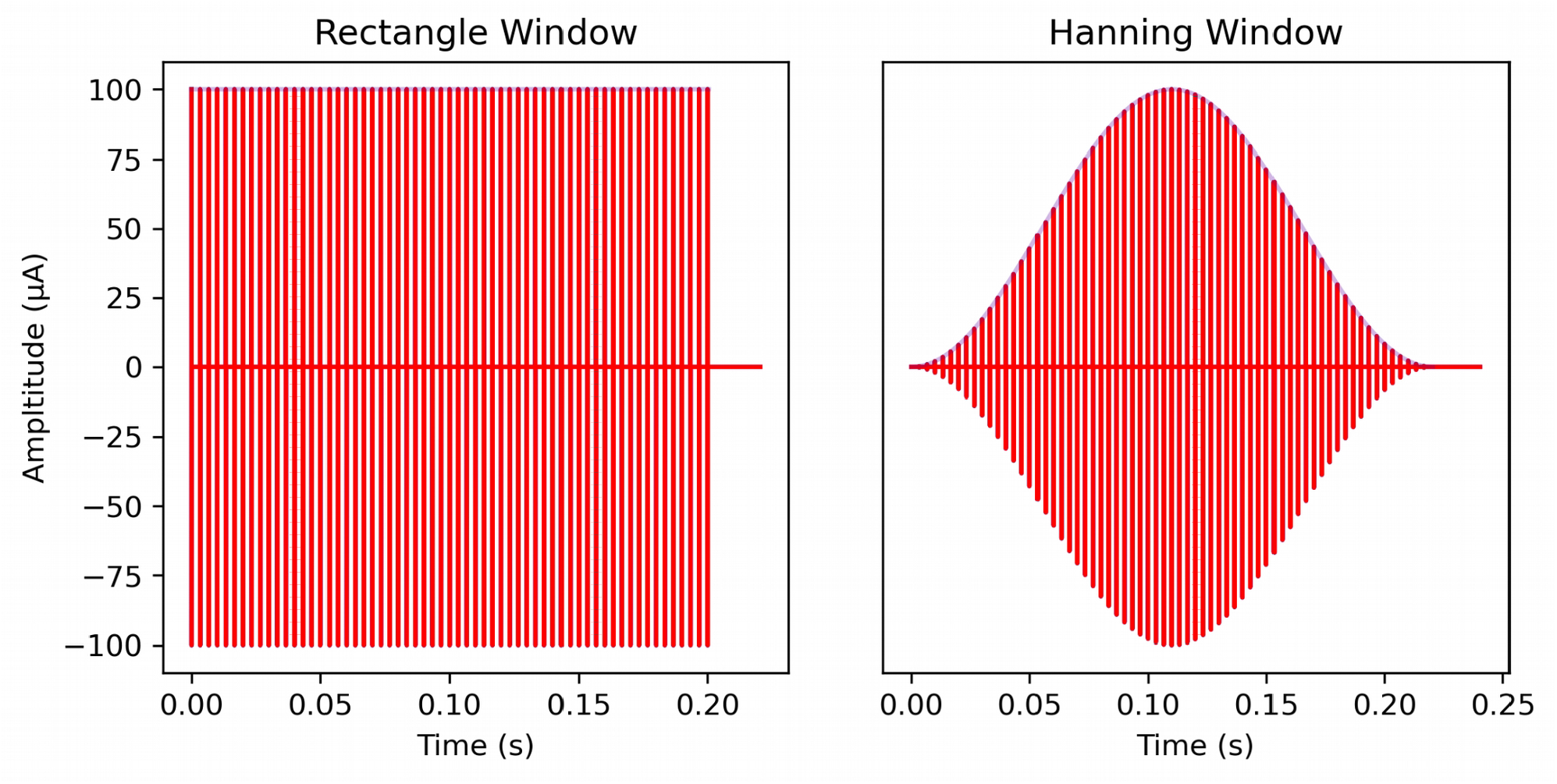
Waveforms of the electrical stimuli for rectangular (left) and Hanning (right) windowed stimulation. Waveforms shown are examples at 300 pps.

If animals struggled with the higher frequencies (particularly 1800 pps) in training sessions, these rates were interleaved with trials at easier pulse rates in order to keep the animals motivated and to allow them to obtain sufficient water rewards for stimuli with both rectangular and Hanning windowed envelopes. All 16 biCI animals were tested for ITD sensitivity under rectangular windowed stimuli at 300 and 900 pps. For rectangular windowed stimuli at pulse rates of 50 and 1800 pps, 14 and 8 CI animals were tested, respectively (Figs. 1, 2, S2). For ITD sensitivity under Hanning windowed envelope, ten CI animals were tested at 50, 300, and 900 pps, and six of them were additionally tested at 1800 pps (Figs. 1, 3, S3).

### Data Analysis

To determine behavioral ITD sensitivity, the proportions of “right” responses (p_R_) an animal made as a function of stimulus ITD were fitted using either linear or cumulative Gaussian sigmoid functions (Figs. 2, 3, S2, S3) as previously described [11]. In brief, psychometric data were fitted with three different, models:

a modified Guassian model

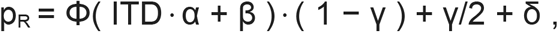

a bounded linear model

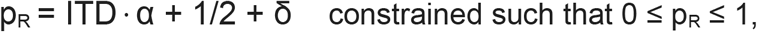

or a null model

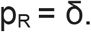

where α captures the animal’s sensitivity to ITD, β captures a possible “ear bias” (that is, 0 ITD may be heard slightly off-center), δ captures a possible spout bias (that is, when guessing, an animal may have an idiosyncratic preference for one side), and γ is a lapse rate, which captures the proportion of times when the animal makes mistakes due to lack of attention or exploratory behavior even though the stimulus should be easy to discriminate. Which of the three possible models provided the best fit to the behavioral data for a given pulse rate and envelope was determined using a deviance test model. The selection criterion is explained in [28]. Once the best fit model was chosen, the slope of the fitted psychometric curve at ITD=0 µs was computed in units of % change in the animal’s preference for choosing the right water spout per µs increase in the value of stimulus ITD. This slope served as a metric for the behavioral sensitivity of the animal to ITD at the given stimulus envelope and pulse rate.

In order to be able to estimate confidence limits and perform statistical inferences on these psychometric slope ITD sensitivity metrics, we then performed a bootstrap resampling analysis. The first step of this analysis consisted of an N-out-of-N resampling of the behavioral data to obtain bootstrap distributions of slope values. For each animal, we collected the N trials the animal performed for a given combination of ITD values, envelope and pulse rate, and drew, with replacement, a bootstrap sample of N trials from that set. Readers versed in probability theory may recognize that this step is equivalent to drawing a random number from a binomial distribution with a “hit” probability equal to the fraction of “right” choices made by the animal for this particular stimulus. After collecting these resampled trials for each ITD tested at the given envelope and pulse rate, the slope of the psychometric function for the resampled data was computed. Repeating this process 1000 times yielded a set of bootstrapped psychometric slopes. These were computed in turn for each animal, envelope type and stimulus pulse rate, and could serve to compute confidence intervals or as empirical distributions which could be further resampled for statistical inference on the effects of changes in stimulus pulse rate or envelope.

To examine the statistical significance of effects in envelope type or pulse rate changes, we decided to carry out 16 pairwise comparisons in total: six comparisons for each pair of pulse rates with rectangular envelopes (see Tab. 1), six with Hanning envelopes (see Tab. 2), and four comparing the two envelope types at each pulse rate tested (see Tab. 3). Because it was not possible to test all animals on every stimulus parameter combination, and because we wished to keep our statistical analysis “within subject”, in order to decide whether changing envelope type or pulse rates had a significant effect, we first identified the subset of animals which were tested at both of the stimulus parameter combinations to be compared. We then carried out resampling of the previously calculated bootstrapped psychometric slopes for each animal to generate mean bootstrap slopes for this cohort. For each animal, one bootstrap slope value was drawn for each of the two stimulus types to be compared, and the values were averaged across the cohort to produce a pair of mean slopes. This process was repeated 16,000 times to generate 16,000 pairs of cohort averaged bootstrap slopes [s_1_, s_2_]_N_, N ∈ {1 … 16,000}. An N of 16,000 repeats was chosen to resolve a smallest p-value of 0.001 after multiple comparison correction (see below). These pairs of resampled, cohort-averaged psychometric slopes were then used to calculate the p-value of the null hypothesis that there were no significant differences between the values s_1_, s_2_ in each pair on average. We first determined whether s_1_ or s_2_ had the smaller mean when averaged over the N cohort resampling trials. Let s_S_ and s_L_ denote the values in each pair that had the smaller (S) or larger (L) mean across N, respectively. Finally, we computed the proportion of the 16,000 pairs for which s_S_ > s_L_. If there was no significant difference in the slopes for each condition, then we would expect s_S_ ≈ s_L_, and s_S_ > s_L_ should then be true almost half the time, but if there was a significant difference, s_S_ > s_L_ should be observed only rarely. The proportion of cases for which s_S_ > s_L_ can therefore serve as an uncorrected p-value for the null hypothesis that the psychometric slopes for each of the two conditions under consideration did not differ significantly. This uncorrected p-value was then multiplied by 16, the total number of comparisons to be performed, and if necessary limited to a maximum of 1, to generate the p-values reported in Tables 1-3. After multiple comparison correction, the smallest p-value that this resampling method can resolve is 0.001, and p-values of 0 returned by this method are therefore reported as <0.001.

The estimated 75% ITD discrimination thresholds were determined from the slopes exhibited by the fitted psychometric curves at 0 µs ITD. As can be seen by inspection of Figures 2, 3, S2, and S3, the psychometrics were either fit directly with a straight-line equation (linear model) or were reasonably well approximated by a straight line tangent over the range of 25-75% right responses (green dashed lines on the sigmoid psychometric functions) with the slope just mentioned. In other words, the psychometric can be reasonably well approximated with a straight line of a constant slope over the range of 25-75% right responses. Consequently, one can estimate the 75% discrimination threshold easily as the change in ITD that would be required to “climb” along the (tangent to the) psychometric function by 25% points, from the 50% level of completely random guessing to the level of 75% correct “right” responses. Note that, due to symmetry, the same absolute change in ITD is required whether one climbs or descends by 25% points along the tangent line. Let *s* denote the slope of the (tangent to the) psychometric in %/μs, and θ_75_ denote the 75% discrimination threshold in μs, then, as just explained, *s*. θ_75_ = 25, or equivalently, θ_75_ = 25/*s*. The mean of the 75% ITD discrimination thresholds, estimated using this simple formula and averaged across animals, are reported in Table 4 for each of the eight different stimulation conditions tested. These 75% thresholds may slightly underestimate thresholds that might be observed in ITD discrimination tasks. First, because they ignore the (usually small) effect of non-linearity for sigmoidal psychometrics, and second, because they estimate the threshold around the animal’s subjective 50% ITD value, rather than at ITD=0 µs. This offsets the possible effects of a small ear-bias which might elevate thresholds in other types of discrimination tasks, depending on the task details. Readers who are interested in precise comparisons of ITD thresholds across studies and species need to keep methodological differences in mind.

## List of abbreviations

biCI: bilateral cochlear implant
CI: cochlear implant
pps: pulses per second
ITD: interaural time difference
ILD: interaural level differences
NH: normal hearing
2-AFC: two-alternative forced choice
ND: neonatally deafened
ABR: auditory brainstem response
eABR: electrically evoked auditory brainstem response
NDCI: neonatally deafened, cochlear implanted
p_R_: proportions of “right” responses
JND: just-noticable difference
N: number of animals

## Declarations

### Ethics approval and consent to participate

Animal experimentation: All procedures involving experimental animals reported here were approved by the Department of Health of Hong Kong (#16-52 to 16-55; 18-81 to 18-83; 20-143 to 20-144 DH/HA&P/8/2/5) or Regierungspräsidium Freiburg (#35-9185.81/ G-17/124), as well as by the appropriate local ethical review committee. All surgery was performed under ketamine and xylazine anesthesia, and every effort was made to minimize suffering.

### Consent for publication

Not applicable.

### Availability of data and materials

The datasets used and analyzed during the current study are available from the corresponding author on reasonable request. All data generated or analyzed during this study are included in this published article and its supplementary information files.

### Authors’ contribution

NRK and JWS: Conceptualization, Resources, Data curation, Software, Formal analysis, Supervision, Funding acquisition, Validation, Investigation, Visualization, Methodology, Writing - original draft, Project administration. ANB: Data curation, Software, Formal analysis, Validation, Investigation, Visualization, Methodology, Writing-original draft. SB: Data curation, Formal analysis, Validation, Investigation, Methodology, Support writing-original draft.

## Acknowledgements

We thank Felix Kleinschroth for his assistance in figure design as well as Felix Kleinschroth, Stella Mayer, and Lakshay Khurana for assisting behavioral training of CI rats.

## Supplementary Materials

**Figure 1 - figure supplement 1 (S1):**
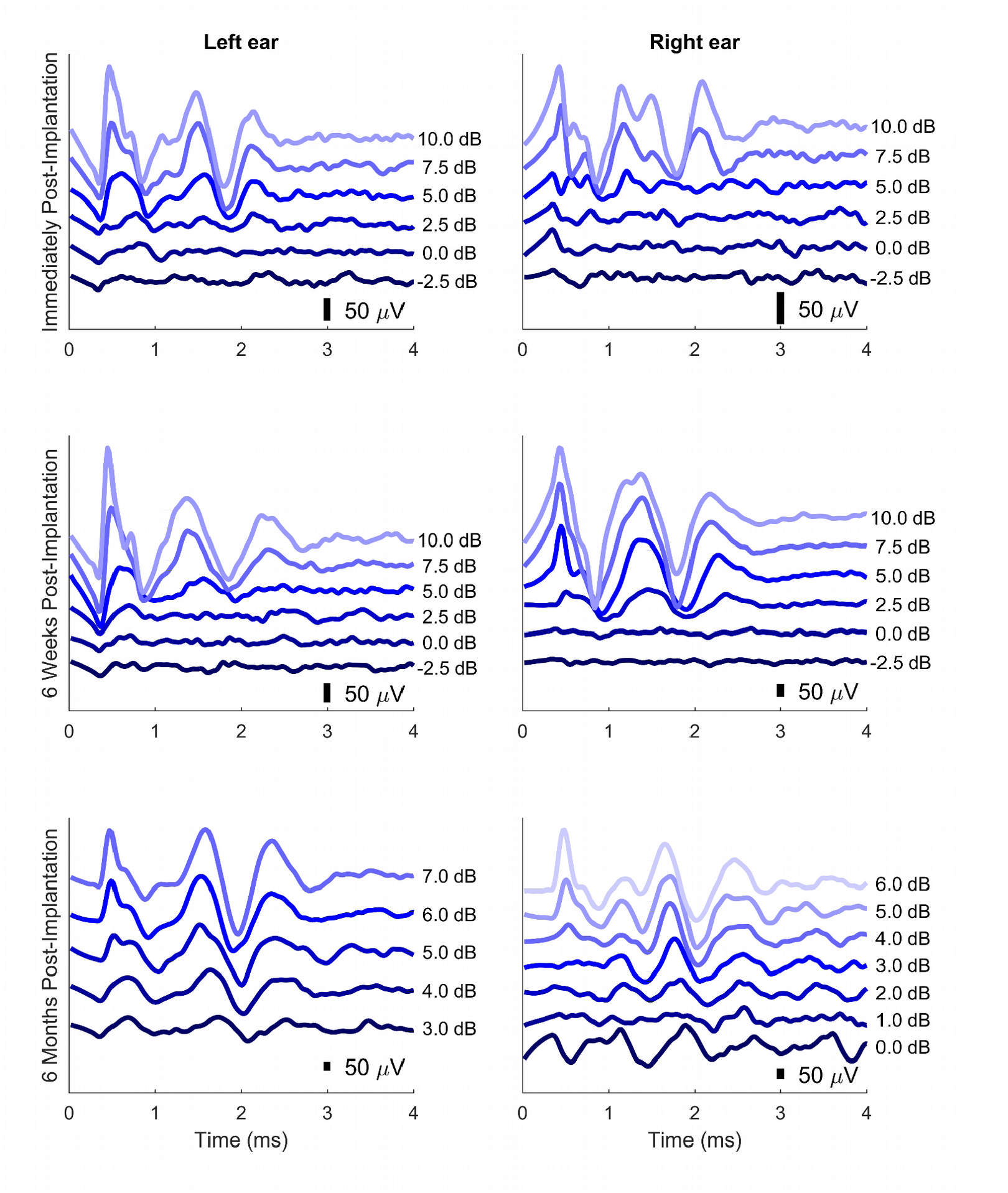
Example eABRs showing immediately after bilateral CI implantation, 6 weeks, and 6 months after implantation thresholds for the left and right ears of a neonatally deafened rat. Scale bars are shown in each plot with reference to 50 μV. Colour represents a different SPL with dark to light colors going from softest to loudest, and 0 dB SPL corresponding to 100 µA current level. Electric artifacts have been removed using interpolation over the duration of the stimulus. For details on the stimulus and presentation, see [11].

**Figure 2 - figure supplement 2 (S2):**
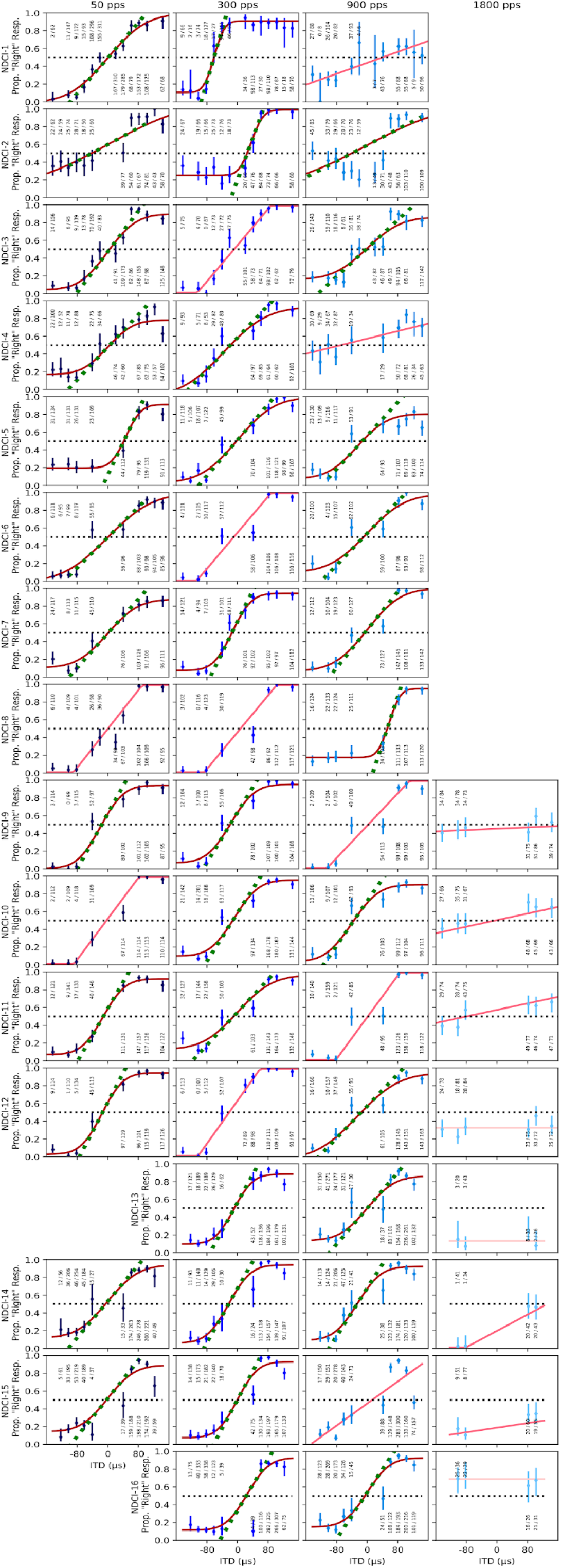
Psychometric functions of all 16 CI rats for rectangular windowed pulse trains at each pulse rate. Each column represents a different pulse rate from 50, 300, 900 and 1800 pulses per second (pps) (left to right). Each row shows the responses for a given animal. The y-axis reflects “right” responses where “right” refers to the right hand spout (Prop. “Right” Resp.). The x-axis shows the tested interaural time difference (ITD) values from – 150 to + 150 µs. Negative ITD values indicate left leading ITDs. Annotations above or below each marker indicate the number of trials the animal chose the right hand side spout over the total number of presentations for a given ITD value. From dark to light the different shades of red indicate sigmoid, linear with bounds or null model psychometric curve fits. Green dashed lines show slopes of psychometric curves at ITD=0 µs. Slopes serve to quantify the behavioral sensitivity of the animal to ITD.

**Figure 3 - figure supplement 3 (S3):**
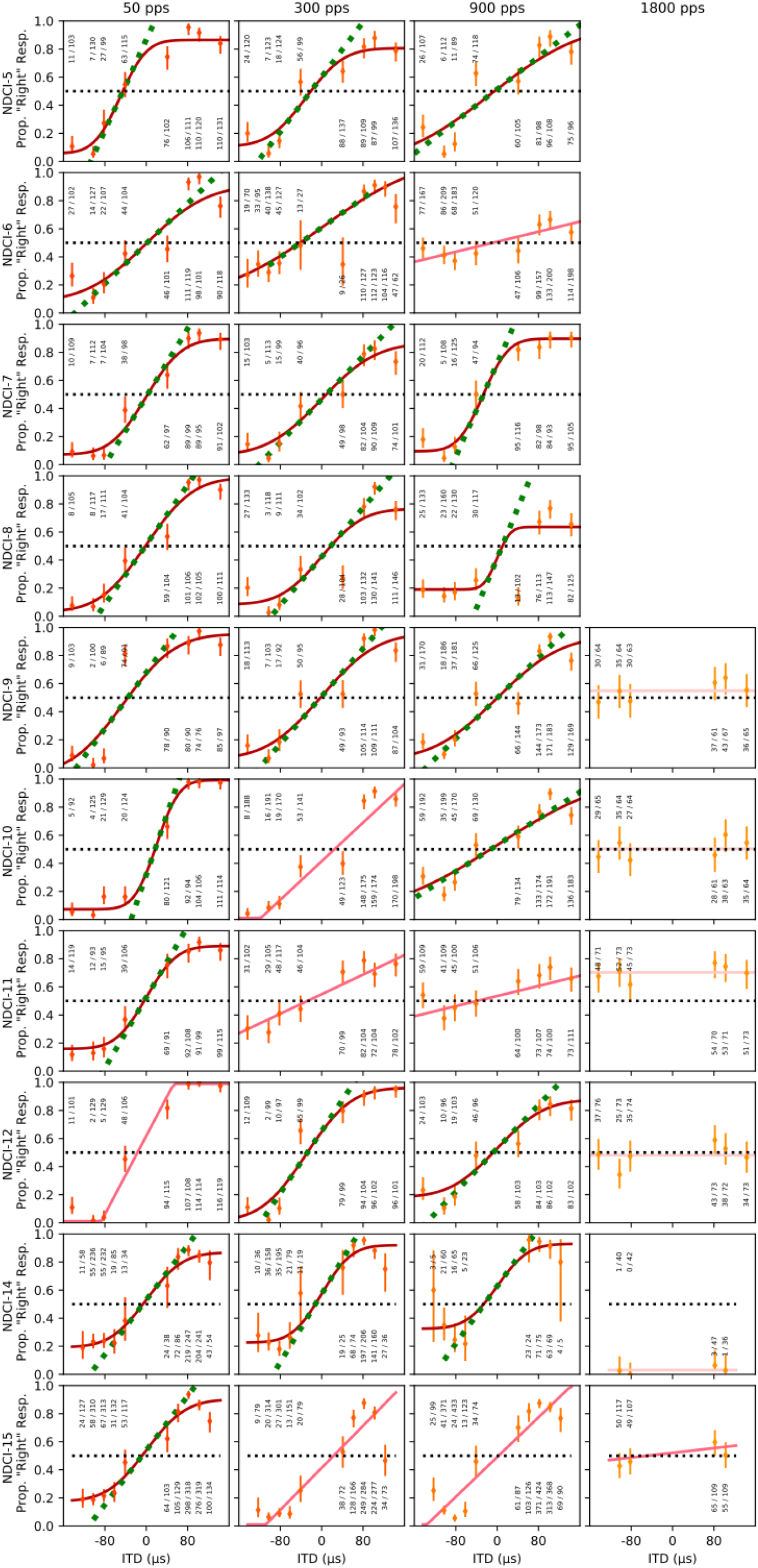
Psychometrics of 10 CI rats for Hanning windowed pulse trains with varying ITDs as in Figure S2. Details are as for S2.

